# Designed mosaic nanoparticles enhance cross-reactive immune responses in mice

**DOI:** 10.1101/2024.02.28.582544

**Authors:** Eric Wang, Alexander A. Cohen, Luis F. Caldera, Jennifer R. Keeffe, Annie V. Rorick, Yusuf M. Aida, Priyanthi N.P. Gnanapragasam, Pamela J. Bjorkman, Arup K. Chakraborty

**Affiliations:** Institute for Medical Engineering and Science, Massachusetts Institute of Technology, Cambridge, MA 02139; Division of Biology and Biological Engineering, California Institute of Technology, Pasadena, CA 91125; Department of Chemical Engineering, Massachusetts Institute of Technology, Cambridge, MA 02139; Department of Physics, Massachusetts Institute of Technology, Cambridge, MA 02139; Department of Chemistry, Massachusetts Institute of Technology, Cambridge, MA 02139; School of Clinical Medicine, University of Cambridge, Hills Rd, Cambridge, CB2 0SP, UK; Ragon Institute of Massachusetts General Hospital, Massachusetts Institute of Technology, and Harvard University, Cambridge, MA 02139

**Keywords:** antibody, computational methods, nanoparticle, protein design, RBD, sarbecovirus, SARS-CoV-2, vaccination

## Abstract

Using computational methods, we designed 60-mer nanoparticles displaying SARS-like betacoronavirus (sarbecovirus) receptor-binding domains (RBDs) by (*i*) creating RBD sequences with 6 mutations in the SARS-COV-2 WA1 RBD that were predicted to retain proper folding and abrogate antibody responses to variable epitopes (mosaic-2_COM_s; mosaic-5_COM_), and (*ii*) selecting 7 natural sarbecovirus RBDs (mosaic-7_COM_). These antigens were compared with mosaic-8b, which elicits cross-reactive antibodies and protects from sarbecovirus challenges in animals. Immunizations in naïve and COVID-19 pre-vaccinated mice revealed that mosaic-7_COM_ elicited higher binding and neutralization titers than mosaic-8b and related antigens. Deep mutational scanning showed that mosaic-7_COM_ targeted conserved RBD epitopes. Mosaic-2_COM_s and mosaic-5_COM_ elicited higher titers than homotypic SARS-CoV-2 Beta RBD-nanoparticles and increased potencies against some SARS-CoV-2 variants than mosaic-7_COM_. However, mosaic-7_COM_ elicited more potent responses against zoonotic sarbecoviruses and highly mutated Omicrons. These results support using mosaic-7_COM_ to protect against highly mutated SARS-CoV-2 variants and zoonotic sarbecoviruses with spillover potential.

## Introduction

Emerging SARS-CoV-2 variants, notably Omicron and its subvariants, have demonstrated the ability to partially evade previous vaccines.^1–9^ While mRNA vaccines have been adapted to include sequences based on existing Omicron strains, they become outdated due to the continuous emergence of new variants.^10^ Moreover, there remains the continuing risk of future pandemics due to spillovers from the pool of existing zoonotic SARS-like betacoronaviruses (sarbecoviruses).^11,12^ Consequently, the development of vaccines capable of safeguarding against future SARS-CoV-2 variants and new viruses derived from sarbecoviruses is critical for public health.

SARS-CoV-2 uses its spike trimer to infiltrate host cells by binding to the host receptor known as angiotensin-converting enzyme 2 (ACE2).^13,14^ Specifically, the receptor-binding domain (RBD) of the spike binds to ACE2, and it can do so only when the RBD adopts an “up” conformation, rather than its usual “down” conformation.^15^ Upon infection or vaccination with the spike trimer, numerous antibodies targeting the RBD are elicited, categorized into four primary types (classes 1, 2, 3, and 4) based on their epitopes (Figure 1A).^15^ The epitopes of class 1 and 2 antibodies typically overlap with the ACE2 binding site on the RBD and have evolved due to immune pressure over time, while class 3 and 4 antibodies bind to more conserved but less accessible (in the case of class 4) epitopes. Notably, class 4 antibodies are sterically occluded even on “up” RBDs, making them challenging to induce using vaccines containing spike trimers. A vaccine capable of eliciting antibodies against the class 4 and class 1/4 (class 4-like antibodies that reach towards the class 1 epitope and sterically occlude ACE2 binding) epitopes^16^ could target conserved sites, providing protection against future SARS-CoV-2 variants and potential sarbecovirus spillovers.

**Figure 1.**
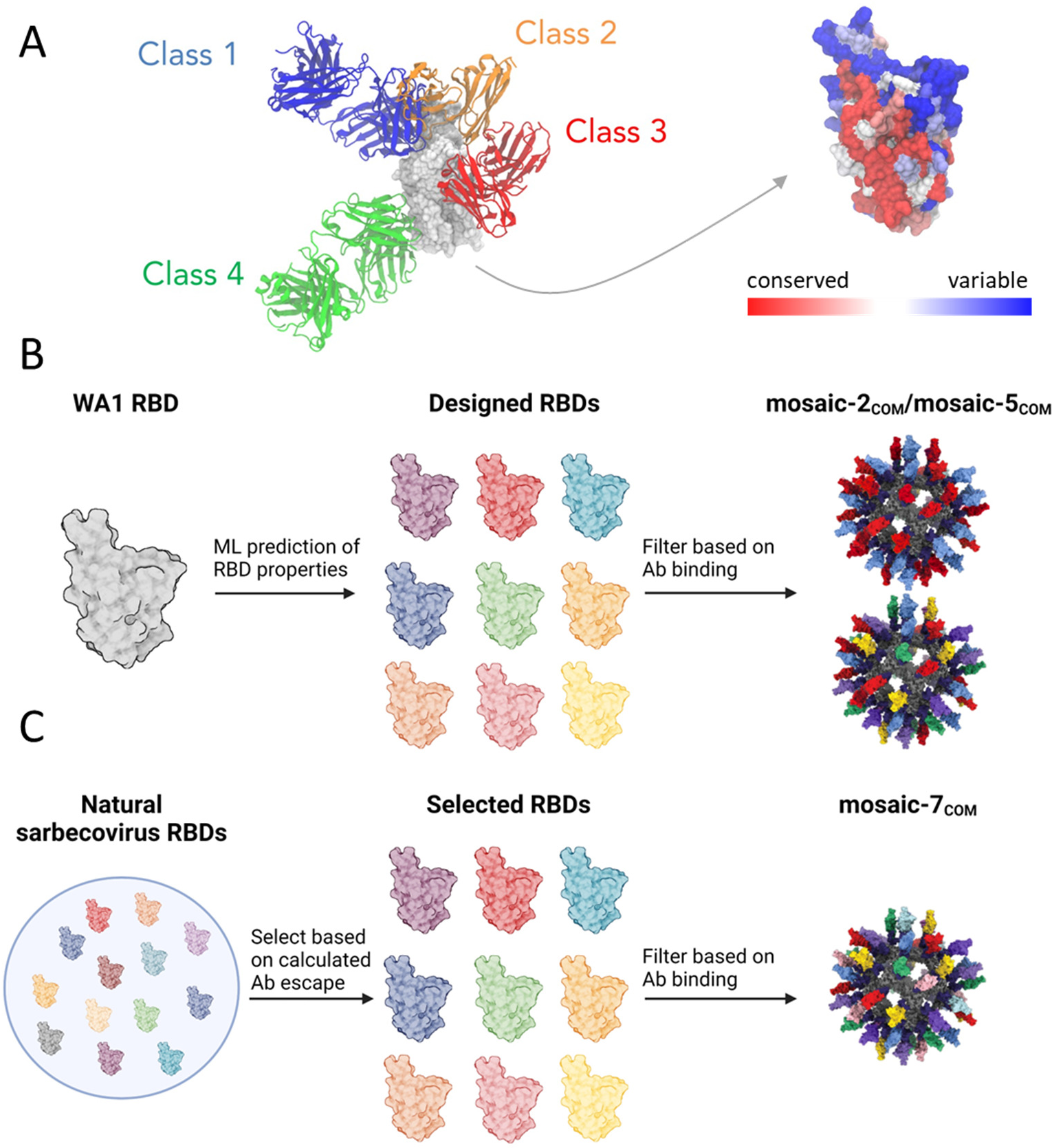
Overview of the design process. (A) Structures of representative class 1 (C102, PDB 7K8M), class 2 (C144, PDB 7K90), class 3 (S309, PDB 7JMX), and class 4 (CR3022, PDB 6W41) antibodies bound to the WA1 SARS-CoV-2 RBD, and the structure of the WA1 RBD (PDB 6W41) colored based on conservation scores calculated using the ConSurf database.^22^ (B) Overview of mosaic-2_COM_ and mosaic-5_COM_ RBD-NP designs. Starting from the WA1 RBD, computational analysis and machine learning models^23^ were used to calculate properties of potential RBD immunogens based on expression, antibody binding, and solubility. A set of selected RBDs were further filtered based on expression and binding measurements and used to construct the mosaic-2_COM_ and mosaic-5_COM_ RBD-NPs. (C) Overview of designing mosaic-7_COM_. A set of 8 RBDs were selected from naturally occurring zoonotic sarbecovirus RBDs to maximize (*i*) sequence diversity and (*ii*) binding to class 3 and 4 but not class 1 and 2 RBD epitopes (RBD epitopes defined as described.^15^ The 8 selected RBDs were further filtered based on experimentally determined properties (see text), and the 7 remaining RBDs were used for mosaic-7_COM_.

Previously, mosaic-8b RBD nanoparticles (RBD-NPs) were developed as a potential pan-sarbecovirus vaccine by using the SpyCatcher-SpyTag system^17,18^ to covalently attach different RBDs with C-terminal SpyTag sequences to a 60-mer mi3 protein NP with N-terminal SpyCatcher proteins in each subunit.^19^ These NPs, which displayed RBDs from SARS-CoV-2 and seven zoonotic sarbecoviruses, were hypothesized to promote the development of cross-reactive antibodies by exposing conserved epitopes and favoring interactions with B cells displaying cross-reactive B cell receptors that can bind bivalently to adjacent conserved regions on the displayed RBDs.^20^ In animal studies, the mosaic-8 RBD-NPs elicited high titers of cross-reactive antibodies^19^ and protected K18-hACE2 transgenic mice^21^ and non-human primates against sarbecovirus challenges^20^. The SpyCatcher-SpyTag system allows various combinations of proteins to be easily attached covalently in various combinations to a SpyCatcher NP, suggesting the intriguing possibility that the displayed RBD sequences could be further optimized to generate NPs that elicit even more potent cross-reactive antibodies.

In this work, we combined computational and experimental approaches to design and test sets of new mosaic RBD-NPs that exhibited improved cross-reactive responses in mice. The first set contained RBDs designed with six mutations relative to the SARS-CoV-2 WA1 strain aimed at maintaining expression and solubility while selectively abrogating antibody binding to class 1 and class 2 RBD epitopes (Figure 1B). The second set contained sarbecovirus RBDs that selectively abrogated class 1 and 2 antibody binding and had the highest sequence diversity among all computationally generated sets (Figure 1C). After experimentally filtering the RBDs for expression, solubility, and antibody binding, we constructed mosaic RBD-NPs and evaluated them in mice. Binding and neutralization titers from naïve mice immunized with RBD-NPs show that our designed RBD-NPs elicited more cross-reactive responses than mosaic-8b and homotypic SARS-CoV-2 Beta RBD-NPs. Deep mutational scanning profiles suggested that the antibody response is focused on class 3 and 4 RBD epitopes for the mosaic-7_COM_ RBD-NP. Finally, serum responses of mice with prior COVID-19 vaccinations showed that mosaic-7_COM_ elicited higher neuralization titers against a range of viral strains compared with mosaic-8b, mosaic-7 (mosaic-8b without SARS-CoV-2 Beta), and bivalent WA1/BA.5 mRNA-LNP. Taken together, these results suggest that designed RBD-NPs, such as mosaic-7_COM_, are promising candidates for potential pan-sarbecovirus vaccines.

## 2 Results

### 2.1 WA1 RBDs were designed to elicit antibodies against less mutated SARS-CoV-2 variants

Our first set of RBD-NPs displayed WA1 RBDs with mutations that were designed to promote generation of cross-reactive antibodies that target relatively conserved epitopes on the RBDs of SARS-CoV-2 variants. Overall, our computational design strategy sought to create RBDs that (*i*) abrogated binding of class 1 and class 2 anti-RBD antibodies but not class 3, 4, and 1/4 antibodies (RBD epitopes defined as described)^15^; (*ii*) were stable and expressed well; and (*iii*) yielded soluble RBD-NPs upon conjugation.

We designed sets of two RBDs to be displayed on a particular NP, with each RBD containing 6 mutations. Although it might be ideal to design RBD-NPs with more variant RBDs, with each containing numerous mutations, introducing many mutations could result in improperly folded RBDs. Our choice of 6 mutations per RBD was informed by our method of predicting relative expression of different RBDs, which is a convolutional neural network trained on deep mutational scanning (DMS) experiments using a library of different RBDs displayed on yeast.^24^ In DMS experiments used to train our neural network, yeast cells displayed RBDs containing random mutations relative to the WA1 strain, and the expression of each variant was measured.^24^ The DMS-generated RBD variants contained between 0 and 7 mutations, so a model trained on these data would not be effective at predicting the expression of variants containing more than 7 mutations. We chose 6 mutations per RBD because this number is below the maximum of 7 mutations and because it is even (we divide the 6 mutations into 3 class 1 escape mutations and 3 class 2 escape mutations).

Previous DMS experiments^25–29^ quantified escape from antibodies (either polyclonal serum antibodies or monoclonal antibodies) in the following way: yeast cells for each RBD mutation were created and sorted into an antibody escape bin based on it not binding to a particular antibody or antiserum. The escape fraction of a RBD mutation is the fraction of yeast cells expressing the mutation that were in the escape bin. An escape fraction of 0 meant that none of the yeast cells expressing the mutation were in the escape bin, while a fraction of 1 meant all yeast cells expressing the mutation were in the bin.

We first considered two RBDs per NP for the following reasons. DMS data on antibody escape at the time we designed the RBD-NPs were evaluated relative to the SARS-CoV-2 WA1 RBD,^25–29^ so it was easiest to design a new RBD with mutations that abrogated binding of WA1-specific class 1 and 2 antibodies. However, new class 1 and 2 anti-RBD antibodies will evolve upon immunization with RBDs that abrogate binding to the usually immunodominant antibodies.^30,31^ Our solution to this problem was to place escape mutations for different RBDs in different positions relative to each other, as the new germline and germinal center (GC) B cells that recognize class 1 and 2 epitopes on one designed RBD would likely not bind bivalently to a second RBD that contains escape mutations in different residues and would thus be at a disadvantage compared to the class 3, 4, and 1/4 antibodies. Based on this hypothesis, one would ideally create an RBD-NP with many different RBDs if their escape mutations were all in different positions. However, we were limited to 6 mutations for the reasons stated above, so we decided to use 2 RBDs per NP to introduce more escape mutations in each RBD and therefore increase the probability of abrogating bivalent antibody binding. However, using only 2 RBDs per NP to create mosaic-2 RBD-NPs results in a higher probability that neighboring RBDs are identical: the average probability of neighboring identical RBDs in a mosaic-2 is 0.5, whereas the average probability of neighboring identical RBDs in a mosaic-8 is 0.125. We also created a mosaic-5_COM_ RBD-NP (Section 2.3) to empirically determine whether displaying more variant RBDs with some shared mutations would result in differences in cross-reactive antibody elicitation.

First, we determined the 20 RBD positions with highest escapes from class 1 and 2 anti-RBD antibodies based on DMS data^25–29^ (Tables S1, S2). We chose to focus on 20 RBD positions because a previous DMS study highlighted ∼20 positions where mutations affected binding to class 1 and 2 antibodies.^27^ Mutations in these 20 positions do not necessarily occur all at once; e.g., the BA.1 SARS-CoV-2 Omicron variant contains substitutions in 15 RBD positions relative to the WA1 RBD. From these 20 positions, we generated 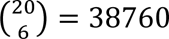 combinations of 6 positions, for both class 1 and class 2 escape positions. Since we have 2 RBDs on a mosaic-2_COM_, we divided each combination of 6 positions into 2 groups of 3, for which there are 10 possible enumerations, generating 387600 sets for class 1 RBD positions and 387600 sets for class 2 positions, as shown in Figure S1. Creating all possible 387600^2^ RBD pairs by combining the class 1 and class 2 sets is computationally infeasible, so we instead randomly sampled ∼800,000 RBD pairs for further evaluation.

Additionally, for a particular escape position, the amino acid mutation with the largest escape fraction that was also not a charged-to-hydrophobic substitution was chosen (Table S2). The decision to avoid charged-to-hydrophobic substitutions was meant to enhance solubility, as preliminary RBD designs showed aggregation when charged-to-hydrophobic mutations were included. For example, RBD residue 484 is a class 2 escape residue,^27^ and the corresponding escape mutation we choose was E484R, which exhibited the largest escape fraction for non-hydrophobic amino acids.^25–29^

The ∼800,000 pairs of RBD sequences were then screened for likelihood of successful expression using a convolutional neural network that was previously trained on DMS data^23^ (Figure 2A,B). We selected RBD pairs for which both RBDs were predicted to express well, which was defined as having a change in expression from WA1 greater than −0.2 log-mean fluorescence intensity (logMFI) based on DMS data.^24^ This threshold was previously chosen such that sequences of circulating variants, which are known to express well because they are found in nature, had predicted logMFI values above this threshold.^24^ Of the ∼800,000 RBD pairs, ∼100,000 were selected that fit the chosen computational expression criterion.

**Figure 2.**
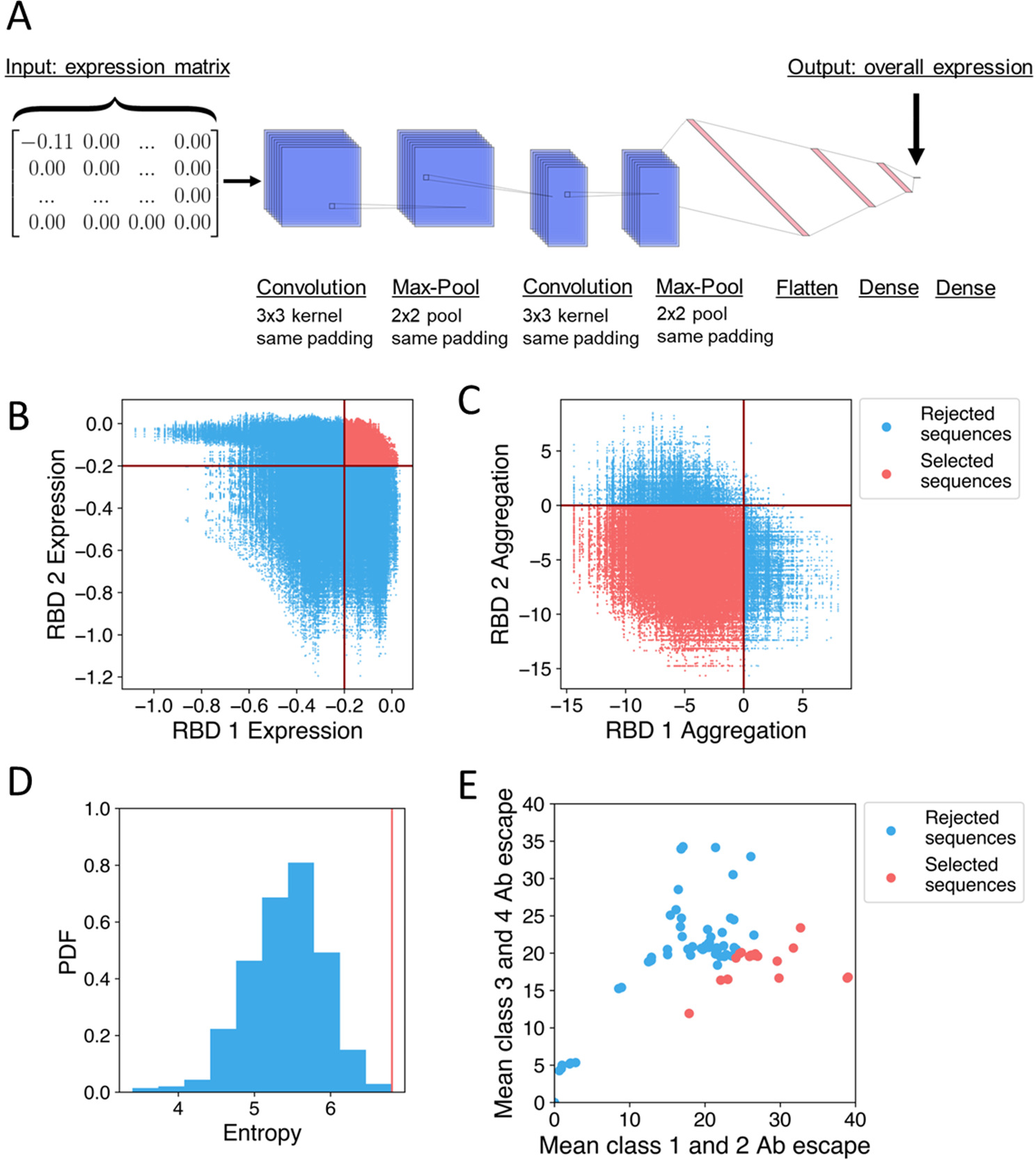
Overview of computational methods. (A) Architecture of the neural network used to predict RBD expression.^23^ The input is an expression matrix, which is the element-wise product (multiplication of entries at the same positions) of the one-hot encoded sequence (each residue is represented as a 20-dimensional vector with entries of 1 for the matching amino acid and 0 for other amino acids) and the matrix of single-mutation expression changes. This is processed through a convolutional neural network to produce the predicted change in expression as an output. (B) ∼800,000 possible RBD sequences are screened for predicted expression relative to the WA1 RBD using a threshold value of −0.2 logMFI. Rejected RBD pairs are in blue and selected pairs are in red. (C) ∼100,000 RBD sequences that passed predicted expression screening and further screened for solubility based on a change in aggregation score relative to WA1 calculated using Aggrescan. Rejected RBD pairs are in blue and selected pairs are in red. (D) The distribution of total mutational entropy over sets of 10 RBDs, and the set selected for experimental testing is the one with maximum entropy indicated by the red line. (E) Mean escape against class 1 and 2 anti-RBD antibodies and the mean escape against class 3 and 4 anti-RBD antibodies for naturally occurring sarbecoviruses. Rejected RBDs are in blue and selected RBDs are in red.

The ∼100,000 selected pairs of RBD sequences were further evaluated for predicted solubility using Aggrescan^32^ to calculate the aggregation score of each RBD in the pair relative to the WA1 RBD (Figure 2C). We selected ∼90,000 RBD pairs for which both RBDs were predicted to be more soluble than the WA1 RBD. The large fraction (∼0.9) of selected pairs suggests that the avoidance of charged-to-hydrophobic mutations in previous steps was effective at preserving predicted solubility.

Of these ∼90,000 RBD pairs, we selected the top 20,000 in terms of total class 1 and class 2 antibody escape (estimated as a sum over the escape fractions for mutated residues on both RBDs) to further reduce recognition of class 1 and class 2 RBD epitopes. We selected 20,000 because the total class 1 and class 2 antibody escape plateaus after the top 20,000 pairs (Figure S2). From these 20,000, we selected a subset for experimental testing. We computationally designed multiple RBD pairs in case some RBDs failed to express, abrogate antibody binding, or remain soluble with limited aggregation. In creating these RBD pairs, we sought to avoid pairs that were very similar. Therefore, we randomly selected sets of 5 RBD pairs, calculated the total mutational entropy, and selected the set with the highest entropy (Figure 2D). More specifically, each set of 5 RBD pairs contained 10 RBDs, and we calculated the Shannon entropy^33^ for each residue over the 10 RBDs. The total mutational entropy was then the sum of the Shannon entropies for all residues. The RBD sequences are reported in Table S3.

### 2.2 Zoonotic sarbecovirus RBDs were selected to elicit cross-reactive antibodies against sarbecoviruses

Additionally, we selected RBDs from various sarbecoviruses to make a new mosaic RBD-NP in a manner distinct from choices for mosaic-8b RBD-NP.^20^ While mosaic-8b used phylogenetics and pandemic potential in its design (selecting clade 1, clade 1b, and clade 2 sarbecovirus RBDs from a study of RBD receptor usage and cell tropism^34^), we instead used antibody binding data to select RBDs. We first obtained a set of 246 non-redundant sarbecovirus RBDs from the NCBI database,^35^ aligned these with the WA1 SARS-CoV-2 RBD using ClustalW,^36,37^ and filtered the alignment for residues 331-531 of the WA1 SARS-CoV-2 spike since these were the WA1 spike residues used for RBD display in DMS experiments.^25–29^ For each RBD in the alignment, we examined its substitutions relative to the WA1 RBD amino acids and calculated the substitutions’ average escapes to class 1, 2, 3, and 4 anti-RBD antibodies from the DMS data. The selective binding B_s_of each RBD was then scored using

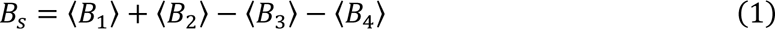

where 〈*B_i_*〉 is the average total escape of an RBD to antibodies of class i. Thus, RBDs that have high escapes from class 1 and 2 antibodies but low escapes from class 3 and 4 antibodies would maximize B_s_. We selected the top 40 sarbecovirus RBDs in terms of *B_s_*. In Figure 2E, we graph the mean class 1 and 2 escapes (〈*B*_l_〉 + 〈*B*_2_〉) and mean class 3 and 4 escapes (〈*B*_3_〉 + 〈*B*_4_〉) for every sarbecovirus RBD and highlight the selected RBDs in red. The selected RBDs clustered towards the lower right region, demonstrating that our calculation of B_s_ selected for high class 1 and 2 escapes and low class 3 and 4 escapes. From the selected RBDs, we generated sets of 8 as in previous studies.^19,20^ We then calculated the fraction of amino acids that were the same for every pair of RBDs (average pairwise amino acid sequence identity as defined to create mosaic-8b). We selected the set of 8 with the lowest average amino acid sequence identity between pairs (Table S4).

### 2.3 Designed RBDs bind class 3 and 4 anti-RBD antibodies and conjugate to form stable RBD-NPs

Before creating mosaic RBD-NPs with the computationally designed RBDs, we experimentally evaluated their expression and binding to characterized anti-RBD monoclonal antibodies, removing any candidates that showed suboptimal properties. First, we expressed the RBDs and purified them from transfected cell supernatants using Ni-NTA affinity chromatography followed by size exclusion chromatography (SEC) (Figure 3A,B). For the designed RBDs, 8 of 10 exhibited expected high levels of expression, while one expressed at low levels (RBD8) and another showed no detectable expression based on SEC chromatograms (RBD3) (Figure 3A). RBD3 and RBD8 were therefore removed from further consideration. Given that 70% of single RBD substitutions eliminated expression in a DMS library,^24^ generating 6-mutant RBDs that preserve expression with an 80% success rate is notably efficient and points to the utility of our neural network predictor. All zoonotic sarbecovirus RBDs expressed effectively (Figure 3B), as expected because well-folded RBDs are likely to be found in nature.

**Figure 3.**
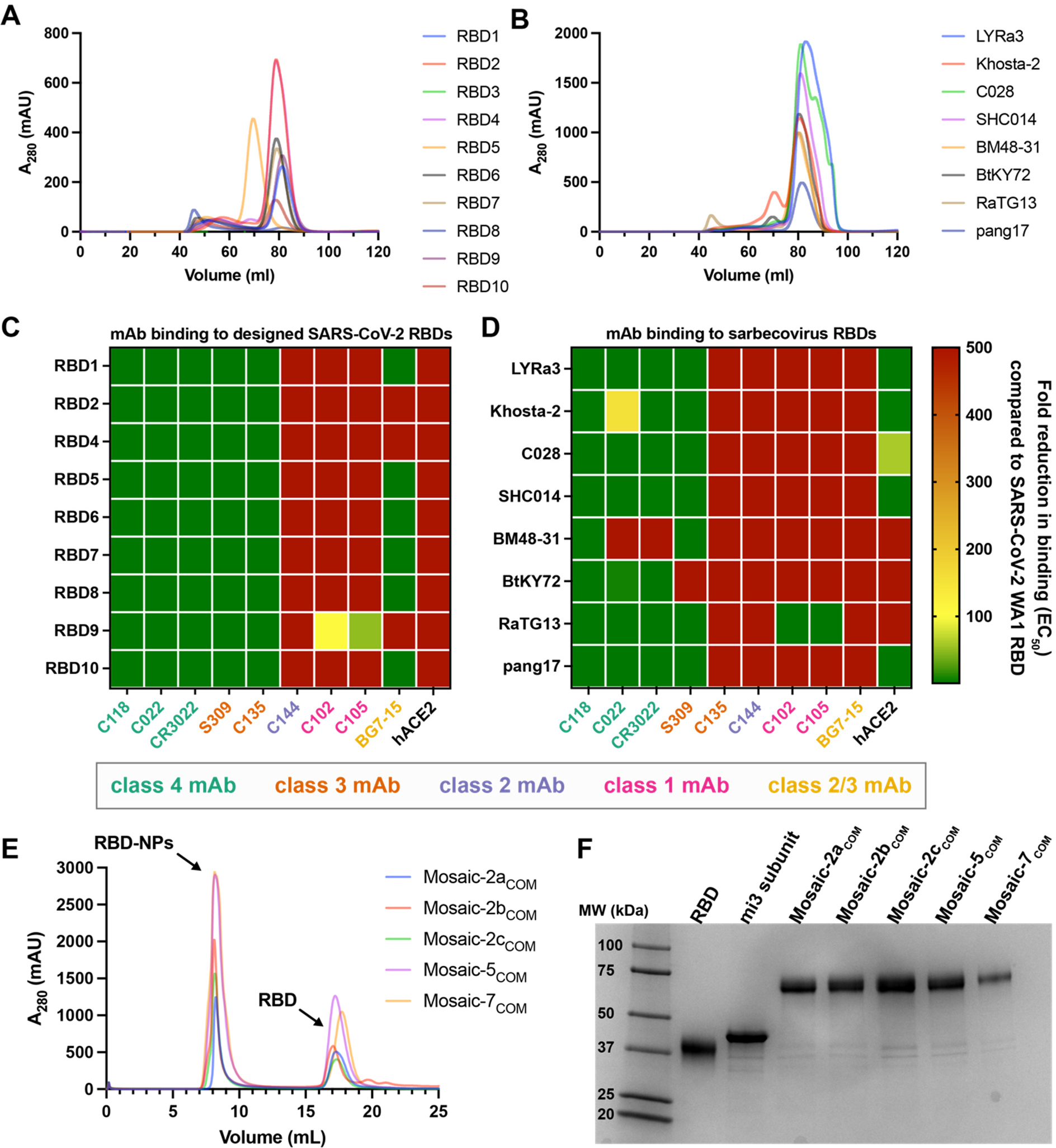
Designed SARS-CoV-2 RBDs and sarbecovirus RBDs exhibit desired properties. (A, B) HiLoad 16/600 Superdex 200 SEC profiles of designed RBDs (A) and sarbecovirus RBDs (B). RBD3 and RBD8 exhibited sub-optimal expression, indicated by no signal for an RBD monomer (RBD3) or a peak in the void volume (RBD8). (C, D) Fold reduction of selected monoclonal anti-RBD antibodies (mAbs) or a human ACE2-Fc construct (hACE2) to designed SARS-CoV-2 RBDs (C) and sarbecovirus RBDs (D) compared with binding to WA1 RBD. (E) Superose 6 Increase 10/300 SEC profiles after SpyTagged RBDs were conjugated to SpyCatcher-mi3 showing peaks for RBD-NPs and free RBDs. (F) SDS-PAGE for each RBD-NP after pooling appropriate SEC fractions.

We then used ELISAs to derive binding EC_50_s of these RBDs to a panel of monoclonal antibodies directed against class 1, 2, 3, and 4 RBD epitopes, demonstrating that the designed RBDs bound class 3 and 4, but not class 1 or class 2, anti-RBD antibodies (Figure 3C). Interestingly, the class 2/3 antibody BG7-15^38^ exhibited mixed results, binding to RBD1, RBD5, RBD6, RBD7, RBD8, and RBD10 but not to RBD2, RBD4, or RBD9 (Figure 3C). Although inconsequential for immunization purposes, none of the designed RBDs bound to a human ACE2-Fc construct because class 1 and 2 escape mutations are located near the ACE2 binding site.^15,25–29^ The EC_50_s of the zoonotic sarbecovirus RBDs for binding the panel of antibodies showed the same trends, although some RBDs (Khosta-2, BM48-31, BtKY72) did not bind all class 3 or 4 antibodies (Figure 3D). RaTG13 RBD retained binding to class 1 antibodies, so it was removed from consideration.

From the designed RBDs, we created 3 RBD-NPs displaying 2 RBDs each (mosaic-2a_COM_, mosaic-2b_COM_, and mosaic-2c_COM_) (Table 1). Although RBD4 and RBD7 showed high expression and bound to class 3 and 4, but not class 1 and class 2, anti-RBD antibodies (Figure 3C), they were not included because they were designed as sets with RBD3 and RBD8, which had been removed. We also created a mosaic-5_COM_ RBD-NP with RBD1, RBD2, RBD4, RBD5, and RBD10 (Table 1) to investigate whether immune responses to an RBD-NP containing more RBDs with some overlapping mutations differed from responses to mosaic-2_COM_ RBD-NPs. RBD6 was excluded because it is similar to RBD10, RBD7 was excluded because it is similar to RBD1, and RBD9 was excluded because it did not completely abrogate binding of class 1 anti-RBD antibodies (Figure 3C). From the zoonotic sarbecovirus RBDs, we used all of the selected RBDs to create a mosaic-7_COM_ RBD-NP, which does not display a SARS-CoV-2 RBD, unlike mosaic-8b RBD-NP (Table 1).^20^

**Table 1.** RBDs in each RBD-NP. RBDs in computationally designed RBD-NPs (mosaic-2_COM_s, mosaic-5_COM_, mosaic-7_COM_) are defined in Tables S3-4. RBDs in mosaic-8b, mosaic-7, and homotypic SARS-CoV-2 are defined in previous studies. ^20,39^

| mosaic-2a<sub>COM</sub> | mosaic-2b<sub>COM</sub> | mosaic-2c<sub>COM</sub> | mosaic-5<sub>COM</sub> | mosaic-7<sub>COM</sub> | mosaic-8b | mosaic-7 | homotypic SARS-CoV-2 |
| --- | --- | --- | --- | --- | --- | --- | --- |
| RBD1 | RBD5 | RBD9 | RBD1 | LYRa3 | Beta |  | Beta |
| RBD2 | RBD6 | RBD10 | RBD2 | Khosta-2 | RaTG13 | RaTG13 |  |
|  |  |  | RBD4 | C028 | Pang17 | Pang17 |  |
|  |  |  | RBD5 | SHC014 | SHC014 | SHC014 |  |
|  |  |  | RBD10 | BM48-31 | WIV1 | WIV1 |  |
|  |  |  |  | BtKY72 | Rs4081 | Rs4081 |  |
|  |  |  |  | Pang17 | Rf1 | Rf1 |  |
|  |  |  |  |  | RmYN02 | RmYN02 |  |

Mosaic-8b, mosaic-7, and homotypic SARS-CoV-2 Beta RBD-NPs were prepared and characterized as described,^19,20,39^ and conjugations to create mosaic-2a_COM_, mosaic-2b_COM_, mosaic-2c_COM_, mosaic-5_COM_, and mosaic-7_COM_ were successful, as demonstrated by SEC (Figure 3E) and SDS-PAGE (Figure 3F).

### 2.4 Designed RBD-NPs elicit cross-reactive binding and neutralization responses in naïve mice

To assess antibody responses to the designed RBD-NPs, we immunized naïve BALB/c mice at days 0, 28, and 56 (Figure 4A). For mosaic-2_COM_ RBD-NP sequential immunizations, mosaic-2a_COM_ was administered on day 0, mosaic-2b_COM_ on day 28, and mosaic-2c_COM_ on day 56. Mice immunized with three doses of mosaic-8b or homotypic SARS-CoV-2 Beta RBD-NPs were included for comparison with other RBD-NPs.

**Figure 4.**
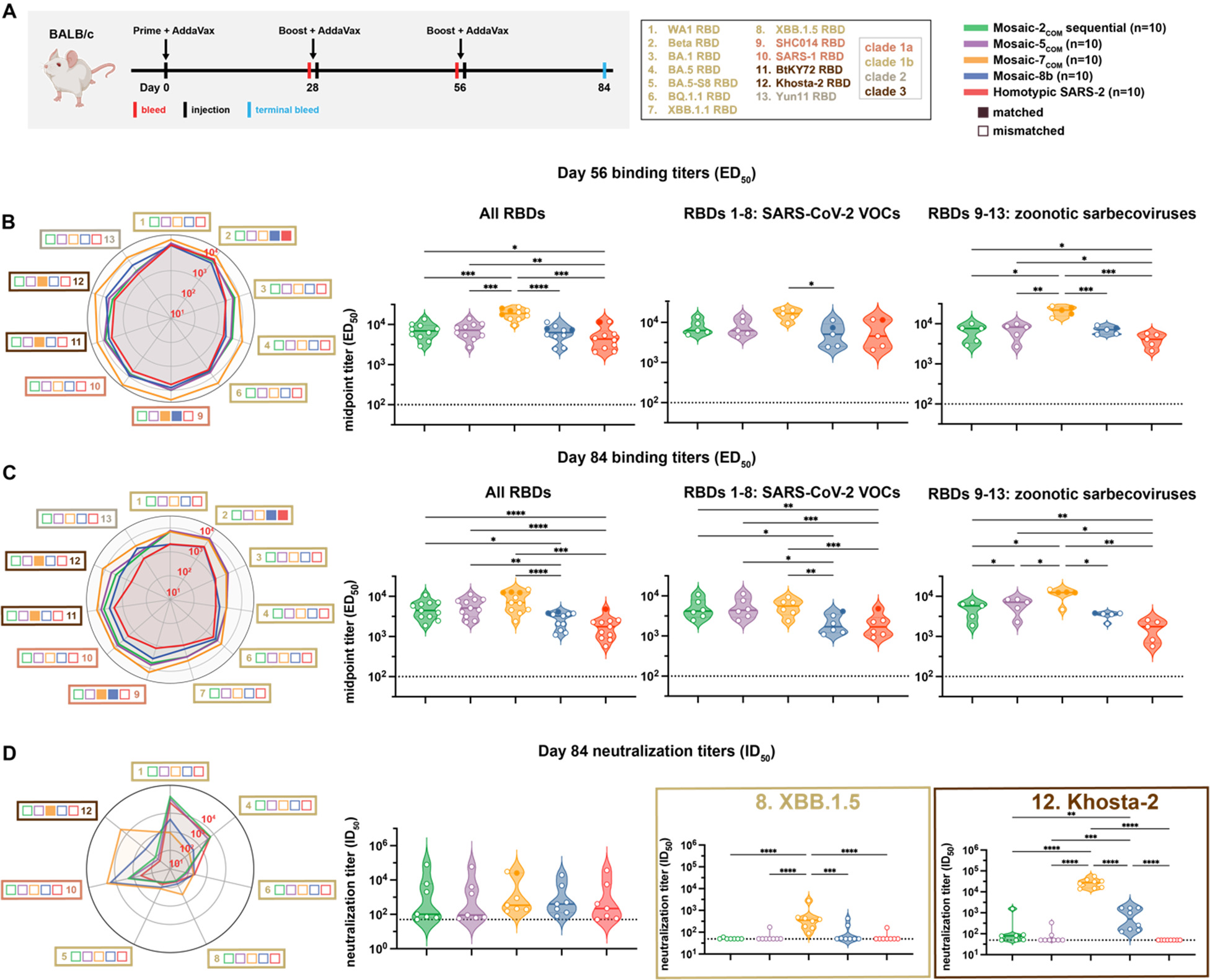
Computationally designed mosaic RBD-NPs elicit cross-reactive antibody binding and neutralization responses in immunized mice. The mean of mean titers is compared in panels B and C by Tukey’s multiple comparison test with the Geisser-Greenhouse correction calculated using GraphPad Prism, with pairings by viral strain. Significant differences between immunized groups linked by horizontal lines are indicated by asterisks: p<0.05 = *, p<0.01 = **, p<0.001 = ***, p<0.0001 = ****. (A) Left: Schematic of immunization regimen. Middle: numbers and colors used for sarbecovirus strains within clades throughout the figure. Right: Colors and symbols (squares) used to identify immunizations (colors) and matched (filled in) versus mismatched (not filled in) viral strains. (B) Left: ELISA binding titers at day 56 for serum IgG binding to RBDs, represented as mean ED_50_ values. Middle left: Means of ELISA binding titers for each immunization. Middle right: Means of ELISA binding titers for each immunization against only SARS-CoV-2 variant RBDs. Right: Means of ELISA binding titers for each immunization against zoonotic sarbecovirus RBDs. Each circle represents the mean serum IgG binding titer against matched (solid circles) and mismatched (open circles) RBDs. (C) Left: ELISA binding titers at day 84 for serum IgG binding to RBDs, represented as mean ED_50_ values. Middle left: Means of ELISA binding titers for each immunization. Middle right: Means of ELISA binding titers for each immunization against only SARS-CoV-2 variant RBDs. Right: Means of ELISA binding titers for each immunization against zoonotic sarbecovirus RBDs. Each circle represents the mean serum IgG binding titer against matched (solid circles) and mismatched (open circles) RBDs. (D) Left: Neutralization titers at day 84 for serum IgG neutralization of pseudoviruses derived from the virus strains in panel A, represented as mean ID_50_ values. Middle left: Means of all neutralization titers for each immunization. Each circle represents the mean neutralization titer against matched (Khosta-2 for mosaic-7_COM_; solid circle) and mismatched (open circles) pseudoviruses. Middle right and right: Neutralization titers against XBB.1.5 and Khosta-2. Each circle represents a neutralization titer from an individual mouse serum sample.

We measured ELISA binding titers against a panel of sarbecovirus RBDs at day 56 and day 84 (Figure 4B,C). Day 56 responses revealed that mosaic-2_COM_ sequential and mosaic-5_COM_ elicited significantly higher titers than homotypic SARS-CoV-2 Beta when comparing means of all RBD titers (Figure 4B, left), as well as when comparing mean binding titers against only zoonotic sarbecovirus strains (Figure 4B, right). Interestingly, mosaic-7_COM_ immunization elicited the highest binding titers against all RBDs including zoonotic sarbecovirus RBDs, rising to significance when comparing mosaic-7_COM_ titers to titers for all other groups.

After three doses of each RBD-NP, the day 84 responses illustrated that the computationally designed RBD-NPs consistently elicited significantly higher binding titers against all evaluated RBDs when compared to mosaic-8b and homotypic SARS-CoV-2 (Figure 4C, left). This was also true when comparing responses against RBDs derived from SARS-CoV-2 VOCs (Figure 4C, middle). However, only the binding responses elicited by mosaic-7_COM_ were significantly better than responses against mosaic-8b when evaluated against zoonotic sarbecovirus RBDs (Figure 4C, right).

Although binding antibody responses showed significant differences between cohorts at day 84, mean neutralization titers across evaluated pseudoviruses, all of which were mismatched except for Khosta-2 (matched for mosaic-7_COM_ but not for the other RBD-NPs), showed no significant differences (Figure 4D). However, mean neutralization titers against individual strains showed some differences (Figure 4D, left). For example, mosaic-7_COM_ elicited lower neutralization titers than mosaic-8b against SARS-CoV-2 WA1 and BA.5, likely because mosaic-7_COM_ does not display a SARS-CoV-2 RBD or an RBD that shares >87% sequence identity with the WA1 or BA.5 RBDs (Figure S3). However, mosaic-7_COM_ elicited significantly higher neutralization titers against XBB.1.5 (mismatched for all RBD-NPs) than the other immunogens (Figure 4D) despite lacking a SARS-CoV-2 RBD. As expected, mosaic-7_COM_ also elicited significantly higher neutralization titers against Khosta-2, a matched strain (Figure 4D). We also found that mosaic-2_COM_ sequential, mosaic-5_COM_, and homotypic SARS-CoV-2 Beta RBD-NPs elicited higher neutralization titers against SARS-CoV-2 WA1 and BA.5 and lower titers against zoonotic sarbecoviruses than the mosaic-7_COM_ and mosaic-8b RBD-NPs, that mosaic-2_COM_ sequential and mosaic-5_COM_ elicited similar neutralization titers as homotypic SARS-CoV-2 Beta RBD-NP against WA1 and BA.5, and that mosaic-2_COM_ sequential and mosaic-5_COM_ elicited higher or similar titers as homotypic SARS-CoV-2 Beta RBD-NP against non-SARS-CoV-2 sarbecoviruses such as SARS-CoV. In addition, binding and neutralizing titers elicited by mosaic-2_COM_ sequential and mosaic-5_COM_ immunizations were generally similar to each other.

It is interesting that the high binding titers against zoonotic sarbecoviruses elicited by mosaic-2_COM_ and mosaic-5_COM_ were not reflected in their neutralization titers, suggesting that mosaic-2_COM_ and mosaic-5_COM_ elicited non-neutralizing anti-RBD antibodies (e.g., against sterically occluded class 4 RBD epitopes^15^) but fewer class 1/4 anti-RBD antibodies, which tend to be more strongly neutralizing^16^.

Taken together, the results suggest that mosaic-2_COM_s, mosaic-5_COM_, and mosaic-7_COM_ would be effective RBD-NPs for eliciting cross-reactive responses in SARS-CoV-2 naïve individuals. In addition, these results validate the approach of using mosaic RBD-NPs composed of computationally designed or selected zoonotic RBDs to elicit broader antibody binding responses to sarbecoviruses.

### 2.5 DMS reveals targeting of conserved RBD epitopes by mosaic RBD-NPs

We further investigated antibody responses raised by mosaic-7_COM_, which elicited both cross-reactive binding and neutralizing titers against sarbecoviruses from different clades (Figure 4C,D). To address which RBD epitopes were recognized, we performed DMS using a SARS-CoV-2 Beta yeast display library^40^ to compare sera from mice immunized with mosaic-7_COM_, mosaic-8b, or homotypic SARS-CoV-2 Beta (Figure 5A). Consistent with a previous DMS comparison of mosaic-8b and homotypic SARS-CoV-2 Beta DMS profiles,^20^ we found stronger DMS profiles for residues within class 3 and 4 RBD epitopes (epitopes defined as described^15^) and weaker DMS profiles in class 2 and class 1 RBD residues for mosaic-7_COM_ and mosaic-8b sera compared to the profile from homotypic SARS-CoV-2 Beta sera. Differences between mosaic-7_COM_ and mosaic-8b sera were difficult to discern across the entire DMS profile but became more apparent when evaluating specific residues on the surface of an RBD (Figure 5B). For example, mosaic-7_COM_ showed higher escape than mosaic-8b at residue 383 (a class 4 residue) and residue 360 (a class 3 residue), suggesting that antibodies recognized an epitope involving those sites. Mosaic-7_COM_ serum also showed little to no escape at RBD residue 484, a class 2 residue that showed high escape from both mosaic-8b and homotypic SARS-CoV-2 Beta sera, suggesting that mosaic-7_COM_ elicited fewer class 2 antibodies against SARS-CoV-2 strains.

**Figure 5.**
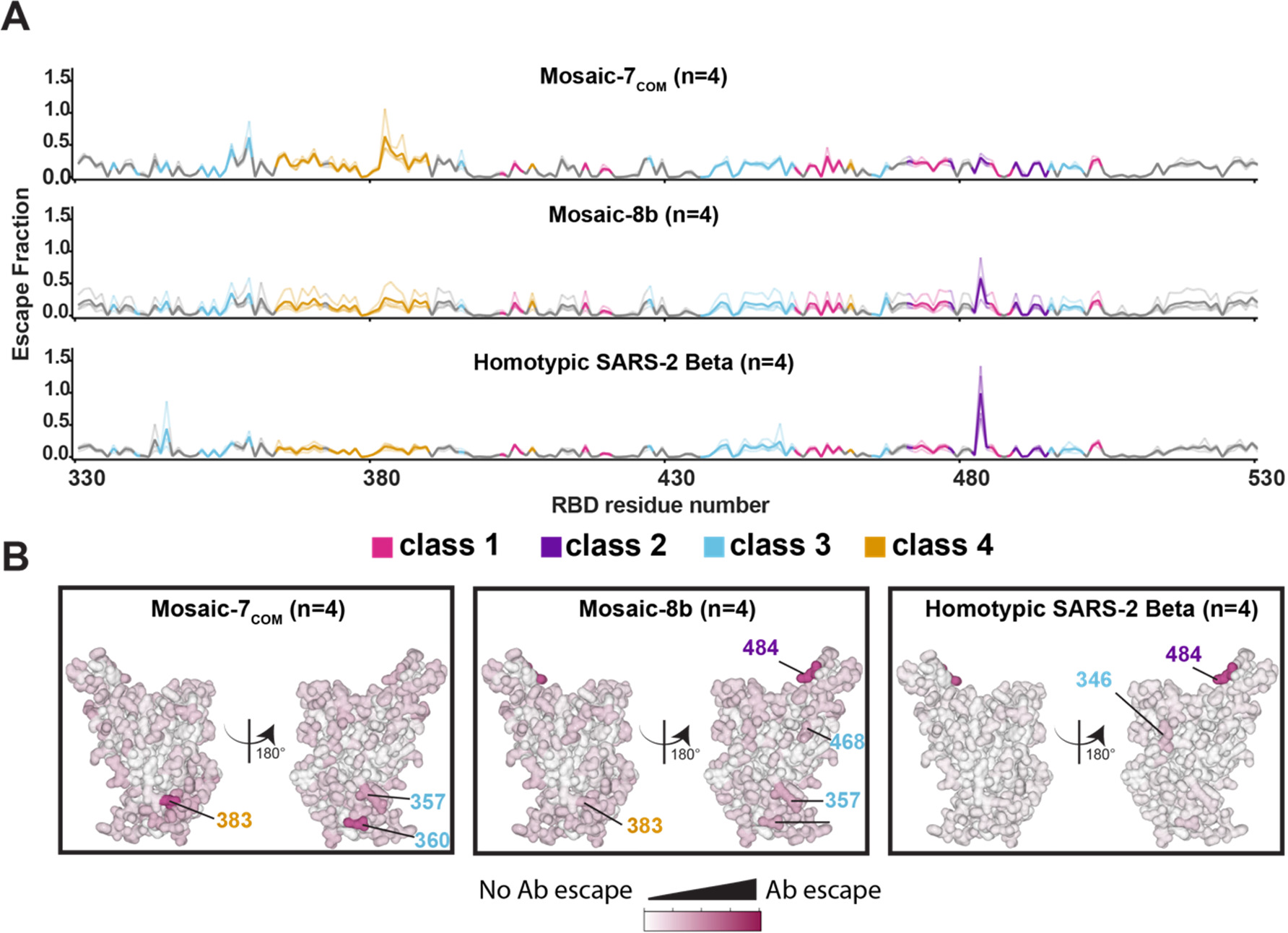
Differences in epitope targeting of antibodies elicited in mice immunized with mosaic and homotypic RBD-NPs. (A) DMS line plots for analyses of sera from mice that were immunized as shown in Figure 4A. DMS was conducted using a SARS-CoV-2 Beta RBD library. The x-axis shows RBD residue positions, and the y-axis shows the total sum of Ab escape for all mutations at a given site, with larger values indicating greater Ab escape. Each faint line represents a single antiserum with heavy lines indicating the average of n=4 sera for each group. Lines are colored differently based on RBD epitopes from the 4 major classes (color definitions are shown in the legend below this panel; gray for residues not assigned to an epitope). (B) Mean site-total antibody escape for a SARS-CoV-2 Beta RBD library determined using sera from mice immunized with the indicated immunogens mapped to the surface of the WA1 RBD (PDB 6M0J). White indicates no escape and dark pink indicates sites with the most escape (residue numbers are denoted with epitope-specific colors as denoted by the legend between panels A and B).

A caveat for interpretation of these DMS results is that the SARS-CoV-2 Beta RBD library was mismatched for mosaic-7_COM_ but matched for mosaic-8b and homotypic SARS-CoV-2 Beta, so there was a greater chance of observing signals in conserved epitopes for mosaic-7_COM_. A matched comparison for both mosaic-7_COM_ and homotypic SARS-CoV-2 Beta could not be made since mosaic-7_COM_ does not display a SARS-CoV-2 Beta RBD. In addition, we previously observed that polyclonal antisera containing antibodies of multiple RBD classes (“polyclass” antibodies) tend to have obscured DMS signals and low escape fractions over all residues compared to DMS signals from monoclonal antibodies or mixtures of anti-RBD antibodies in which one antibody class is dominant.^39^ Thus, it is possible that the differences between DMS profiles for mosaic-7_COM_ and mosaic-8b were dampened by the polyclass nature of the elicited antibodies against the mosaic RBD-NPs.

### 2.6 Mosaic-7COM elicited superior cross-reactive responses in mice with prior COVID-19 vaccinations

We next investigated the impact of prior COVID-19 vaccinations on mosaic-7_COM_ by immunizing BALB/c mice that had previously been vaccinated with two doses of a WA1 Pfizer-equivalent mRNA-LNP vaccine followed by a bivalent WA1/BA.5 mRNA-LNP vaccine (Figure 6A). We immunized mice with two doses of mosaic RBD-NPs (either mosaic-7_COM_, mosaic-8b, or mosaic-7, mosaic-8b without SARS-CoV-2 Beta RBD^39^), or an additional dose of bivalent WA1/BA.5 mRNA-LNP. Results for the mosaic-8b and mosaic-7 cohorts in this experiment were previously described^39^; here, we compare those results to mosaic-7_COM_ immunizations because both mosaic-7 RBD-NPs lack a SARS-CoV-2 RBD, whereas mosaic-8b includes the SARS-CoV-2 Beta RBD (Table 1). As previously discussed, levels of binding antibodies after animals had received the same course of mRNA-LNP vaccines showed significant differences in titers elicited by the pre-vaccinations across cohorts^39^ (Figure S4B-C, day 0). We therefore used baseline corrections (see Methods) to account for different mean responses at day 0 in each of the groups for the data shown in Figure 6. (Binding data without baseline corrections are shown in Figure S4B-C.) Neutralization potencies at day 0 were similar for all cohorts and therefore were not baseline-corrected.^39^

**Figure 6.**
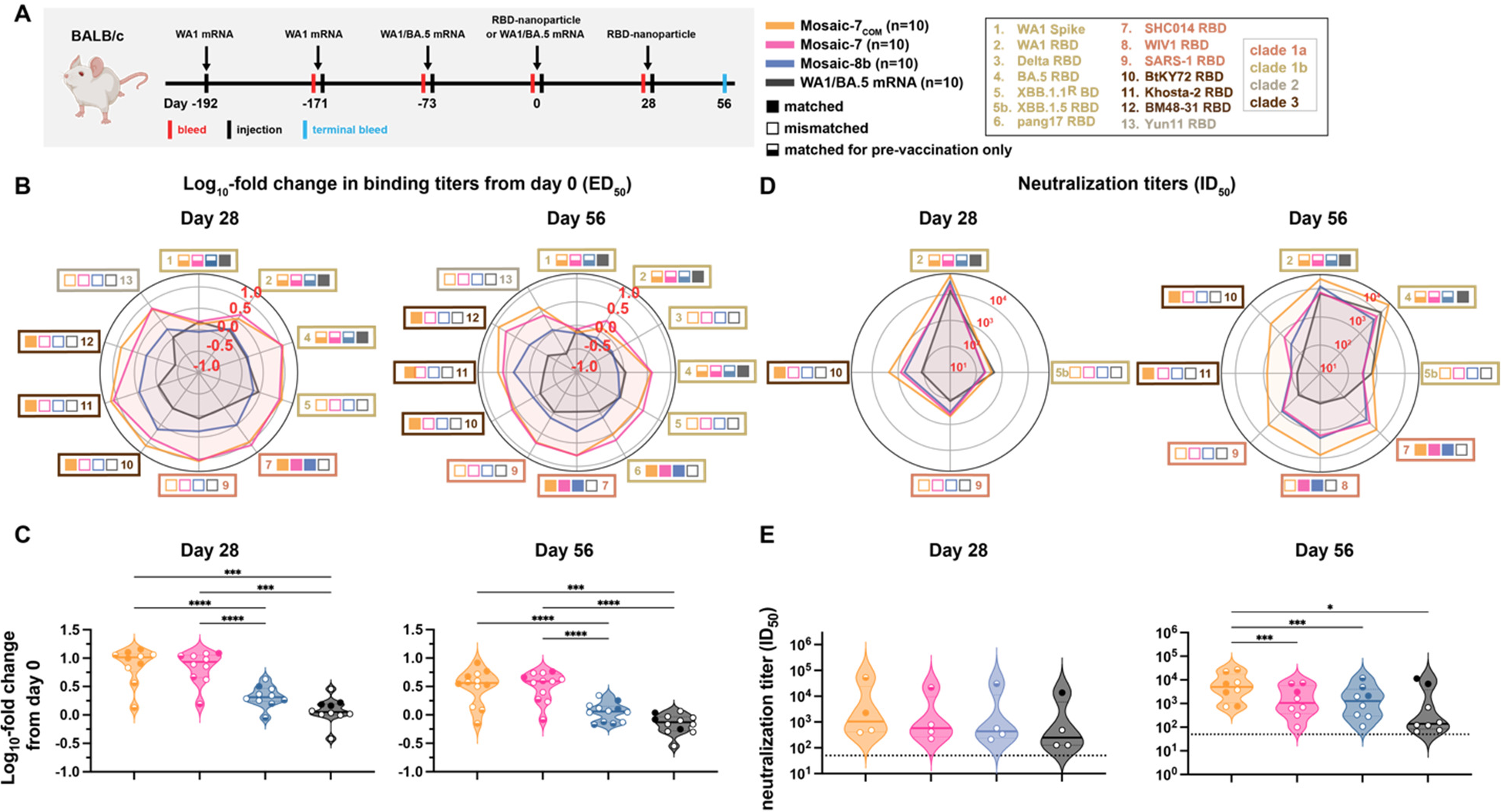
Mosaic-7_COM_ immunization in pre-vaccinated mice elicits superior cross-reactive antibody responses. The mean of mean titers is compared in panels C and E by Tukey’s multiple comparison test with the Geisser-Greenhouse correction calculated using GraphPad Prism, with pairings by viral strain. Significant differences between immunized groups linked by horizontal lines are indicated by asterisks: p<0.05 = *, p<0.01 = **, p<0.001 = ***, p<0.0001 = ****. Binding responses at day 0 (before NP or other vaccine immunizations) showed significant differences across cohorts in titers elicited by the pre-vaccinations.^39^ To account for different mean responses at day 0 between cohorts, we applied baseline corrections (see Methods). Uncorrected binding data for panels B and C are shown in Figure S4B-C. (A) Left: Schematic of vaccination regimen. Mice were pre-vaccinated with mRNA-LNP encoding WA1 spike and bivalent WA1/BA.5 prior to prime and boost immunizations with RBD-NPs at day 0 and day 28 or an additional WA1/BA.5 mRNA-LNP immunization at day 0. Middle: Colors and symbols (squares) used to identify immunizations (colors) and matched (filled in), mismatched (not filled in), or matched to pre-vaccination (half-filled in) viral strains (squares). Right: numbers and colors used for sarbecovirus strains within clades throughout the figure. (B) Log_10_ mean fold change in ELISA ED_50_ binding titers from day 0 at the indicated days after priming with the indicated immunogens against spike or RBD proteins from the indicated sarbecovirus strains (numbers and color coding as in panel A). (C) Log_10_ means of fold change in ELISA titers for each type of immunization at the indicated days. Each circle represents the log_10_ mean fold change in ED_50_ titers from mice against a single viral strain of sera from mice that were immunized with the specified immunogen (solid circles=matched; open circles=mismatched; colors for different strains defined in panel A). (D) Mean change in neutralization ID_50_ titers from day 0 at the indicated days against the indicated sarbecovirus strains (numbers and color coding as in panel A). (E) Means of all neutralization titers for each type of immunization at the indicated days. Each circle represents the mean neutralization IC_50_ titer against a single viral strain of sera from mice that were immunized with the specified immunogen (solid circles=matched; open circles=mismatched; colors for different strains defined in panel A).

Day 28 and 56 log_10_ fold changes in ELISA binding titers (after prime and boosting RBD-NP immunizations) are shown in Figure 6B. At both timepoints, RBD-NPs generally more strongly boosted binding titers than a second dose of a bivalent WA1/BA.5 mRNA-LNP vaccine, especially against zoonotic sarbecoviruses. Mosaic-8b boosted titers less than mosaic-7 and mosaic-7_COM_ against all viral strains, and mosaic-7 and mosaic-7_COM_ largely boosted titers to similar extents. Mean log_10_ fold changes in binding titers (Figure 6C) showed that mosaic-7 and mosaic-7_COM_ both boosted binding titers significantly better than mosaic-8b or WA1/BA.5 mRNA-LNP.

Mosaic-7 and mosaic-8b elicited similar neutralization titers at both day 28 and 56 (Figure 6D), whereas mosaic-7_COM_ elicited higher neutralization titers than other immunogens against both zoonotic sarbecoviruses and SARS-CoV-2 variants at day 56. Notably, mosaic-7_COM_ antisera neutralized XBB.1.5 with equal potency as antisera from mice boosted with a second dose of WA1/BA.5 mRNA-LNP, whereas antisera from mosaic-7 and mosaic-8b prime/boosted animals exhibited lower potencies, resulting in statistically significant differences in mean neutralization titers between mosaic-7_COM_ and other immunogens (Figure 6E).

Although the mean of means of baseline-corrected binding titers for mosaic-7_COM_ and mosaic-7 cohorts were similar to each other (Figure 6B) and both were significantly higher than titers for mosaic-8b (Figure 6C), the non-baseline corrected binding titers for these groups at day 28 and day 56 showed differences (Figure S4B): e.g., mosaic-7_COM_ sera elicited significantly higher mean of mean titers than mosaic-7 or mosaic-8b (Figure S4C). The higher titers for mosaic-7_COM_ compared with mosaic-7 could be related to the fact that the mosaic-7_COM_ cohort started with higher antibody binding titers than mosaic-7 at day 0 (prior to RBD-NP immunizations), but this does not apply to differences with the mosaic-8b cohort since the mosaic-7_COM_ and mosaic-8b titers at day 0 were equivalent (Figure S4C).

Taking all experimental results into account, mosaic-7_COM_ showed broader and more potent binding and neutralization than either of its mosaic NP counterparts, suggesting that mosaic-7_COM_ more efficiently elicits broader binding and more potently neutralizing antibodies in a pre-vaccinated animal model.

## 3 Discussion

Multivalent NPs have emerged as a useful platform for developing new vaccines against mutable pathogens including influenza,^41,42^ HIV,^43–45^ RSV,^44^ and SARS-CoV-2.^19,20,39,46–54^ Many of these are homotypic SARS-CoV-2 spike- or RBD-NPs, which display multiple copies of only one SARS-CoV-2 spike or RBD and are analogous to homotypic SARS-CoV-2 Beta RBD-NP, which we used for previous and current comparisons with mosaic-8b RBD-NPs.^20,39^ Displaying different variants on a single NP could provide broader protection, as previous studies demonstrated that the mosaic-8b RBD-NP protected K18-hACE2 transgenic mice from a mismatched SARS-CoV challenge whereas homotypic SARS-CoV-2 Beta RBD-NP did not.^20^ Recently, we also studied the impact of prior COVID-19 vaccinations on mosaic-8b RBD-NP vaccines, finding that mosaic-8b still elicited cross-reactive antibodies, primarily by boosting them from prior vaccinations.^39^

The “plug-and-display” SpyCatcher-SpyTag system^17,18^ has been used with a variety of antigens^19,20,42,55–57^ and can be easily adapted to make mosaic NPs with different antigenic compositions. This flexibility led us to evaluate whether designing RBD sequences using computational methods and available data could enhance elicited cross-reactive antibody responses beyond those of previously-studied RBD-NPs such as mosaic-8b, mosaic-7, or homotypic SARS-CoV-2 Beta.^39^ We designed two sets of RBD-NPs that are effective for different targets. First, we combined DMS data, machine learning, and structure-based solubility predictions to design RBDs with 6 mutations relative to the WA1 RBD, which were then displayed as sets of 2 RBDs on NPs. RBDs within a set were designed to reduce bivalent B cell receptor binding to class 1 and 2 RBD epitopes while maintaining bivalent binding to class 3 and 4 epitopes, and these designed RBDs were then used to create the mosaic-2_COM_s and mosaic-5_COM_ RBD-NPs. Since these RBDs only included a few substitutions relative to the WA1 RBD, the mosaic-2_COM_s and mosaic-5_COM_ were designed to be effective against less mutated SARS-CoV-2 variants. Second, we used DMS escape measurements^25–29^ and sequence diversity to select naturally occurring sarbecovirus RBDs to create the mosaic-7_COM_ RBD-NP, which was designed to be effective against zoonotic sarbecoviruses and heavily mutated SARS-CoV-2 variants. Binding and neutralization titers following immunization of naïve mice suggested that the designed RBD-NPs are indeed most effective against their respective proposed targets, and all of the computationally designed RBD-NPs were superior to previously described RBD-NPs (homotypic SARS-CoV-2 Beta for less mutated SARS-CoV-2 variants and mosaic-8b/mosaic-7 for zoonotic sarbecoviruses and heavily mutated SARS-CoV-2 variants.^39^

Although less mutated SARS-CoV-2 variants are no longer circulating in humans, our computational method for generating mosaic-2_COM_s and mosaic-5_COM_ could still be useful. For example, the mosaic-2_COM_s and mosaic-5_COM_ were just as potent as homotypic SARS-CoV-2 Beta RBD-NP against less mutated SARS-CoV-2 variants and more potent against zoonotic sarbecoviruses and heavily mutated SARS-CoV-2 VOCs. Thus, the mosaic-2_COM_s and mosaic-5_COM_ exhibited superior properties compared with homotypic Beta RBD-NP. Our designs had an 80% success rate of producing folded SpyTagged RBDs that expressed well, suggesting that these methods could be used for other SARS-CoV-2 variants or for other viruses.

To address the ongoing rise of SARS-CoV-2 Omicron variants and potential zoonotic sarbecovirus spillovers, we introduce mosaic-7_COM_ as an RBD-NP that provides more effective cross-reactive responses than other mosaic RBD-NPs (mosaic-8b and mosaic-7)^20,39^ in both naïve and pre-vaccinated mice. DMS results, although possibly obscured by the presence of multiple antibody classes in polyclass sera,^39^ suggested that mosaic-7_COM_ elicited more antibodies binding to class 3 and 4 epitopes than mosaic-8b and homotypic SARS-CoV-2 Beta RBD-NPs. Additionally, mosaic-7_COM_ elicited fewer antibodies recognizing an epitope involving the class 2 RBD residue 484, suggesting that mosaic-7_COM_ effectively redirects the antibody response from variable epitopes to conserved epitopes compared to mosaic-8b. We note that both of the mosaic-RBD NPs that lack a SARS-CoV-2 RBD (mosaic-7_COM_ and mosaic-7) outperformed mosaic-8b in mice that had received mRNA-LNP vaccines expressing SARS-CoV-2 WA1 and Omicron BA.5 spikes, consistent with a lack of expansion of SARS-CoV-2 specific immunodominant antibodies that target variable epitopes in mice receiving RBD-NPs that did not include a SARS-CoV-2 RBD. Additionally, mosaic-7_COM_ and mosaic-7 outperform mosaic-8b despite the possibility that pre-existing high affinity antibodies could block variable SARS-CoV-2 epitopes and enhance cross-reactive responses to mosaic-8b.^30,58–62^ Although removal of a SARS-CoV-2 RBD from mosaic-8b to create mosaic-7 improved binding responses in pre-immunized mice, mosaic-7_COM_ elicited significantly higher neutralizing titers than mosaic-7 against both zoonotic sarbecoviruses and SARS-CoV-2 variants, supporting its use as a pan-sarbecovirus vaccine in populations that have been exposed to SARS-CoV-2.

A guiding principle for creating effective mosaic RBD-NPs is maximizing the diversity of RBDs displayed on a single NP to focus the response on conserved epitopes.^20,63^ This can be done by increasing the number of variant RBDs to decrease the probability that B cell receptors can crosslink between immunodominant variable epitopes on adjacent identical RBDs. In this work, the enhanced cross-reactive responses elicited by mosaic-7_COM_ illustrate that computationally optimizing protein sequences is another way to increase displayed RBD diversity. Overall, our results support the integration of computational methods with vaccine design and the further evaluation of designed RBD-NPs, particularly mosaic-7_COM_, as potential pan-sarbecovirus vaccines.

## 4 Methods

### 4.1 Resource availability

#### 4.1.1 Lead contact

Further information and requests for resources should be directed to and will be fulfilled by the lead contact, Arup K. Chakraborty.

#### 4.1.2 Materials availability

All unique/stable reagents generated in this study will be made available on request by the lead contact with a completed materials transfer agreement.

#### 4.1.3 Data and code availability

All original code and data files for the computational designs have been deposited at https://github.com/ericzwang/designed_mosaic_NPs and are publicly available. DMS data will be posted in GitHub upon publication.

### 4.2 Methods details

#### Protein expression and purification

Monoclonal IgGs, a soluble SARS-CoV-2 trimer with 6P stabilizing mutations, and a human ACE2-Fc construct^64^ were produced as previously described.^16,19,20^

Vectors encoding the protein sequences for computationally designed RBDs were assembled using Gibson cloning from an insert encoding residues 319-541 of the SARS-CoV-2 WA1 RBD with the indicated substitutions (Table 1; Table S3). For the selected sarbecovirus RBDs in Table S4, RBDs were also assembled using Gibson cloning from inserts encoding the indicated residues. RBDs used for mosaic-8b, mosaic-7, and homotypic SARS-2 were expressed as described previously.^39^ Sarbecovirus RBDs from SARS-CoV-2 Beta (GenBank QUT64557.1), SARS-CoV-2 WA1 (GenBank MN985325.1), SARS-CoV-2 BA.5-S8,^65^ LYRa3 (AHX37569.1),^65^ Khosta-2 CoV (QVN46569.1), SHC014-CoV (GenBank KC881005), BM48-31-CoV (GenBank NC014470), BtKY72-CoV (GenBank KY352407), Yun11-CoV (GenBank JX993988), WIV1-CoV (GenBank KF367457), RaTG13-CoV (GenBank QHR63300), SARS-CoV (GenBank AAP13441.1), Rs4081-CoV (GenBank KY417143), RmYN02-CoV (GSAID EPI_ISL_412977), Rf1-CoV (GenBank DQ412042), and pangolin17-CoV (GenBank QIA48632) were constructed as previously described.^15,19,20,39^ Briefly, RBDs used for conjugations were encoded with a C-terminal hexahistidine tag (6xHis; G-HHHHHH) and SpyTag003 (RGVPHIVMVDAYKRYK)^66^ for conjugating onto SpyCatcher003-mi3 to form mosaic NPs. RBDs used for ELISAs were encoded with a C-terminal Avi tag (GLNDIFEAQKIEWHE) followed by a hexahistidine tag (6xHis; G-HHHHHH).

RBDs were expressed and subsequently purified via His-tag affinity and SEC purification from transiently-transfected Expi293F (ThermoScientific) supernatants.^20^ RBDs for immunizations in mRNA-LNP pre-vaccinated mice were prepared as described for the analogous experiment.^39^

#### Preparation of RBD-NPs

SpyCatcher003-mi3 nanoparticles for the RBD-NPs were expressed and purified as described.^39^ NPs were aliquoted and flash frozen in liquid nitrogen before being stored at −80 °C until use.

For immunizations in naïve mice, SpyTagged RBDs were conjugated onto SpyCatcher003-mi3 as described.^13,14^ Briefly, equimolar amounts of RBDs for each mosaic NP were mixed before addition of purified SpyCatcher003-mi3, with the final concentration having 2-fold molar excess of total RBD to mi3 subunit: an equimolar mixture of 2 RBDs for the mosaic-2_COM_s, 5 RBDs for mosaic-5_COM_, 7 RBDs for mosaic-7_COM_ and mosaic-7, 8 RBDs for mosaic-8b, or only SARS-CoV-2 Beta for homotypic RBD-NPs. Reactions were incubated overnight at room temperature in Tris-buffered saline (TBS) on an orbital shaker. Free RBDs were purified the next day by SEC on a Superose 6 10/300 column (GE Healthcare) and equilibrated with PBS (20 mM sodium phosphate pH 7.5, 150 mM NaCl). RBD-NP conjugations were assessed by SDS-PAGE. Concentrations of conjugated mi3 nanoparticles are reported based on RBD content, determined using a Bio-Rad Protein Assay. RBD-NPs were aliquoted and flash frozen in liquid nitrogen before being stored at −80 °C until use.

RBD-NPs for immunizations in mRNA-LNP pre-vaccinated mice were prepared as described for the analogous experiment.^39^

#### Mice

6- to 7-week-old female BALB/c mice (Charles River Laboratories) were housed at Labcorp Drug Development, Denver, PA for immunizations. All animals were healthy after being weighed and monitored for 7 days preceding the start of the study. Mice were randomly assigned to experimental groups of 10. Cages were kept in a climate-controlled room at 68-79 °C at 50 ± 20% relative humidity. Mice were provided Rodent Diet #5001 (Purina Lab Diet) ad libitum. Mouse procedures were approved by the Labcorp Institutional Animal Care and Use Committee.

#### Immunizations

For immunizations in naïve animals, RBD-NPs were diluted using Dulbecco’s PBS and mixed 1:1 (v/v) with AddaVax prior to immunization, for a final vaccine dose of 5 ug of RBD equivalents in 0.1 mL total volume. RBD-NP immunizations were administered intramuscularly (IM) via both right and left hindleg (50 µl each). Mice were immunized three times at days 0, 28, and 56, bled via tail vain at days 0, 28, and 56, with a terminal bleed via cardiac puncture at day 84. Blood samples were allowed to clot, and sera were collected and stored at −80°C prior to use.

As previously described,^39^ mice used in the in the pre-vaccination study were vaccinated IM with 20 µL of WA1 mRNA-LNP at days −192 and −171 containing 1 µg mRNA diluted in PBS and 20 µL of WA1/BA.5 mRNA-LNP (0.5 µg WA1 and 0.5 µg BA.5 mRNA) at day −73. Mice were then immunized IM with 5 µg of protein nanoparticle (RBD equivalents) in 100 µL containing 50% v/v AddaVax adjuvant on days 0 and 28 or received an additional dose of 1 µg WA1/BA.5 mRNA-LNP at day 0. Mice were bled and sera were collected as described above.

#### Reagents used for pre-vaccinations

Pfizer-equivalent mRNA-LNP formulations for WA1 and BA.5 were purchased from Helix Biotech as described.^39^ Bivalent WA1/BA.5 mRNA LNP was prepared by mixing WA1 mRNA-LNP and BA.5 mRNA-LNP 1:1 by mRNA mass.

#### Antibody binding and neutralization assays

Binding of characterized anti-RBD monoclonal antibodies and a human ACE2-Fc construct to RBDs was assessed as described.^16^ Monoclonal antibodies were assigned to RBD epitopes based on structural studies as described.^15^

Binding to purified RBDs or spike proteins was assessed using serum samples from immunized mice by ELISA as described.^39^ We used Graphpad Prism 10.1.1 to plot and analyze binding curves, assuming a one-site binding model with a Hill coefficient to obtain midpoint titers (ED_50_ values for serum ELISAs, EC_50_ values for monoclonal antibody ELISAs). ED_50_/EC_50_ values were normalized and mean ED_50_/EC_50_ values were calculated as described.^39^ For pre-vaccinated mouse data shown in Figure 6, ED_50_s were normalized by dividing the ED_50_ response at day 28 and day 56 (after RBD-NP immunizations or an additional mRNA-LNP immunization) over the ED_50_ response at day 0 to account for differences in binding responses between groups in the pre-vaccination cohorts (Figure S4C; left panel). Figure S4 compares non-baseline correcting binding responses across cohorts (panels B and C) with baseline corrected binding responses (panels D and E).

Lentiviral-based pseudoviruses were prepared and neutralization assays conducted and luciferase activity was measured as relative luminescence units (RLUs) as described.^39^ Relative RLUs were normalized to RLUs from cells infected with pseudotyped virus in the absence of antiserum. Half-maximal inhibitory dilutions (ID_50_ values) were derived using 4-parameter nonlinear regression in Antibody Database.^67^

#### DMS

DMS studies used to map epitopes recognized by serum Abs were performed in biological duplicates using independent SARS-CoV-2 Beta-based mutant libraries (generously provided by Tyler Starr, University of Utah) as described previously.^40^ Serum samples were heat inactivated and depleted for yeast binding as described before.^39^ DMS was performed and escape fractions were calculated and analyzed as described.^39,68^ Raw data will be available in a GitHub repository upon publication.

Static line plot visualizations of escape maps were created using Swift DMS as described.^39,68^ Line heights in static line plot visualizations of escape maps (created as described^39,68^) indicate the escape score for that amino acid mutation. RBD epitopes were classified using previously-described class 1, 2, 3, and 4 nomenclature.^15^ For structural visualizations, an RBD surface of PDB 6M0J was colored by the site-wise escape metric at each escape site, with dark pink scaled to be the maximum escape fraction used to scale the y-axis for serum Abs and white indicating no escape. Residues that exhibited the greatest escape fractions were marked with their residue number and colored according to RBD epitope class.

## 5 Acknowledgments

This work was supported by the National Science Foundation Graduate Research Fellowship: 1745302 (E.W.), the National Institutes of Health: 1-R61-AI161805-01 and U19AI057229 (A.K.C.), the National Institutes of Health: P01-AI165075 (P.J.B.), Wellcome Leap (P.J.B.), Bill and Melinda Gates Foundation: INV-034638 (P.J.B.), the Coalition for Epidemic Preparedness Innovations (CEPI) (P.J.B.), and the Merkin Institute for Translational Research (Caltech). E.W. and A.K.C. acknowledge the MIT SuperCloud and Lincoln Laboratory Supercomputing Center for providing HPC resources that have contributed to the research results reported within this work.

We thank Jesse Bloom (Fred Hutchinson), Allie Greaney (University of Washington), and Tyler Starr (University of Utah) for RBD libraries and help setting up DMS at Caltech, Noor Youssef (Harvard Medical School) and Ziyan Wu (Caltech) for radar plot Python scripts, Jost Vielmetter, Luisa Segovia, Annie Lam, and the Caltech Beckman Institute Protein Expression Center for protein production, Igor Antoshechkin and the Caltech Millard and Muriel Jacobs Genetics and Genomics Laboratory for Illumina sequencing, Chengcheng Fan for generating RBD-NP models, Labcorp Drug Development (Denver, PA) for performing mouse immunizations, and Anthony West for calculating the probabilities of identical neighboring RBDs.

## 6 Author Contributions

Conceptualization: E.W, A.K.C., A.A.C., P.J.B.; Methodology: E.W, A.K.C. (computations); J.R.K., A.V.R., Y.M.A., P.N.P.G. (experiments); Computation and Software: E.W., A.K.C.; Investigation: E.W. (computations); J.R.K., A.V.R., Y.M.A., P.N.P.G. (experiments); Writing – original draft: E.W., A.K.C, A.A.C, L.F.C., P.J.B.; Writing – review and editing: E.W., A.K.C, A.A.C, L.F.C., P.J.B.; Visualization: E.W., A.A.C, L.F.C.; Supervision: A.K.C, P.J.B.; Project Administration: A.K.C., P.J.B.; Funding: A.K.C., P.J.B.

## 7 Competing Interests

A.K.C. is a consultant (titled “Academic Partner”) for Flagship Pioneering, consultant and Strategic Oversight Board Member of its affiliated company, Apriori Bio, and is a consultant and Scientific Advisory Board Member of another affiliated company, Metaphore Bio.

P.J.B. and A.A.C. are inventors on a US patent application (17/523,813) filed by the California Institute of Technology that covers mosaic RBD-NPs. P.J.B. is a scientific advisor for Vaccine Company, Inc. and for Vir Biotechnology.

