## Supplementary Information for "Designed mosaic nanoparticles enhance cross-reactive immune responses in mice"

**Table S1**. Antibodies in the deep mutational scanning data[1-5] used to calculate escape.

| Antibody | RBD epitope class[6] |
|---|---|
| COV2-2196 | class 1 |
| S2H14 | class 1 |
| COV2-2165 | class 1 |
| COV2-2832 | class 1 |
| S2E12 | class 1 |
| LY-CoV016 | class 1 |
| REGN10933 | class 1 |
| C105 | class 1 |
| S2X58 | class 2 |
| S2D106 | class 2 |
| S2H13 | class 2 |
| S2H58 | class 2 |
| S2X16 | class 2 |
| LY-CoV555 | class 2 |
| C121 | class 2 |
| C144 | class 2 |
| C002 | class 2 |
| COV2-2096 | class 2 |
| COV2-2050 | class 2 |
| COV2-2479 | class 2 |
| C135 | class 3 |
| S2X227 | class 3 |
| COV2-2499 | class 3 |
| REGN10987 | class 3 |
| S309 | class 3 |
| COV2-2130 | class 3 |
| C110 | class 3 |
| CR3022 | class 4 |
| COV2-2677 | class 4 |
| S2X35 | class 4 |
| COV2-2094 | class 4 |
| S304 | class 4 |
| S2H97 | class 4 |
| COV2-2082 | class 4 |
| S2X259 | class 4 |

**Table S2**. The 20 RBD positions with highest escapes from class 1 and 2 anti-RBD antibodies[6] based on DMS data[1-5] and their assigned amino acid. An assigned amino acid was selected as the amino acid with the largest mean escape fraction relative to the WA1 amino acid, which also was not a charged-to-hydrophobic mutation.

| Antibody RBD-epitope class | RBD position with high escape | Assigned amino acid |
|---|---|---|
| 1 | 417 | Y |
| 1 | 420 | Q |
| 1 | 449 | D |
| 1 | 455 | R |
| 1 | 456 | G |
| 1 | 460 | L |
| 1 | 472 | P |
| 1 | 473 | F |
| 1 | 475 | N |
| 1 | 476 | S |
| 1 | 484 | K |
| 1 | 485 | S |
| 1 | 486 | A |
| 1 | 487 | K |
| 1 | 489 | L |
| 1 | 496 | V |
| 1 | 498 | D |
| 1 | 500 | Q |
| 1 | 501 | A |
| 1 | 504 | L |
| 2 | 346 | D |
| 2 | 447 | D |
| 2 | 449 | K |
| 2 | 450 | T |
| 2 | 452 | R |
| 2 | 455 | G |
| 2 | 456 | A |
| 2 | 472 | P |
| 2 | 473 | F |
| 2 | 475 | N |
| 2 | 481 | K |
| 2 | 483 | R |
| 2 | 484 | R |
| 2 | 485 | K |
| 2 | 486 | P |
| 2 | 487 | K |
| 2 | 489 | V |
| 2 | 490 | K |
| 2 | 493 | K |
| 2 | 494 | R |

**Table S3**. Designed RBDs with their specific mutations relative to the SARS-CoV-2 WA1 RBD. 3 mutations are in RBD positions with high escape against class 1 antibodies[6] based on DMS data[1-5] (class 1 escape mutations), and 3 mutations are in RBD positions with high escape against class 2 antibodies (class 2 escape mutations). RBDs that pass experimental validation in terms of expression (Figure 3B) and binding to class 3 and 4 antibodies but not class 1 and 2 antibodies (Figure 3D) are bolded.

| RBD Pair | RBD ID | Class 1 escape mutations | Class 2 escape mutations |
|---|---|---|---|
| 1 | **RBD1** | L455R, F486A, N487K | E484R, Y489V, F490K |
| | **RBD2** | K417Y, A475N, Q498D | G447D, L452R, Q493K |
| 2 | RBD3 | L455R, F456G, N487K | E484R, F490K, Q493K |
| | **RBD4** | A475N, Y489L, N501A | Y449K, L452R, F486P |
| 3 | **RBD5** | K417Y, F486A, Q498D | F456A, E484R, S494R |
| | **RBD6** | L455R, A475N, N487K | I472P, F490K, Q493K |
| 4 | **RBD7** | A475N, F486A, N487K | L452R, E484R, Y489V |
| | RBD8 | L455R, F456G, Y473F | G485K, F490K, Q493K |
| 5 | **RBD9** | F486A, N487K, Y489L | Y449K, L452R, E484R |
| | **RBD10** | L455R, I472P, A475N | F456A, F490K, S494R |

**Table S4**. Selected sarbecovirus RBDs along with their GenBank[7] accession numbers, chosen residue numbers based on alignment with the SARS-CoV-2 WA1 RBD, and clade. RBDs that passed experimental validation in terms of expression (Figure 3B) and binding to class 3 and 4 antibodies but not class 1 and 2 antibodies (Figure 3D) are bolded. The clade is defined as described in Starr et al.[8]

| Virus | Accession | Residue number | Clade |
|---|---|---|---|
| **LYRa3** | AHX37569.1 | 310-527 | 1a |
| **Khosta-2** | QVN46569.1 | 307-522 | 3 |
| **C028** | AAV98001.1 | 306-523 | 1a |
| **SHC014** | QJE50589.1 | 307-524 | 1a |
| **BM48-31** | YP 003858584.1 | 310-524 | 3 |
| **BtKY72** | APO40579.1 | 309-526 | 3 |
| **pang17** | QIQ54048.1 | 317-549 | 1b |
| RaTG13 | QHR63300.2 | 319-541 | 1b |

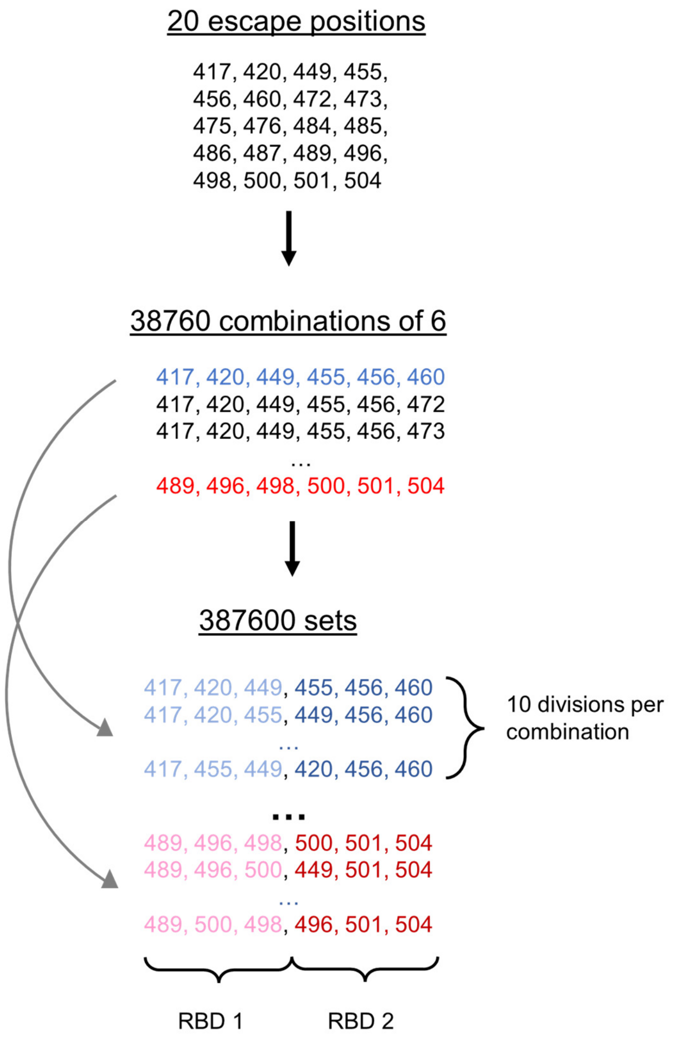


**Figure S1.** Illustration of how sequences for class 1 escape mutations were generated. From the 20 RBD positions with highest escapes against class 1 anti-RBD antibodies, we generated all 38760 possible combinations of 6 positions. For each combination, we further generated all possible ways to divide the 6 positions into 2 groups of 3, of which there were 10 possible divisions, resulting in 387600 sets of positions. For each set, 1 group of 3 positions was assigned to RBD1 (colored as light blue or pink), and the other group of 3 positions was assigned to RBD2 (colored as dark blue or red). RBD1 and RBD2 would be mutated in their assigned positions to the amino acids listed in Table S2.

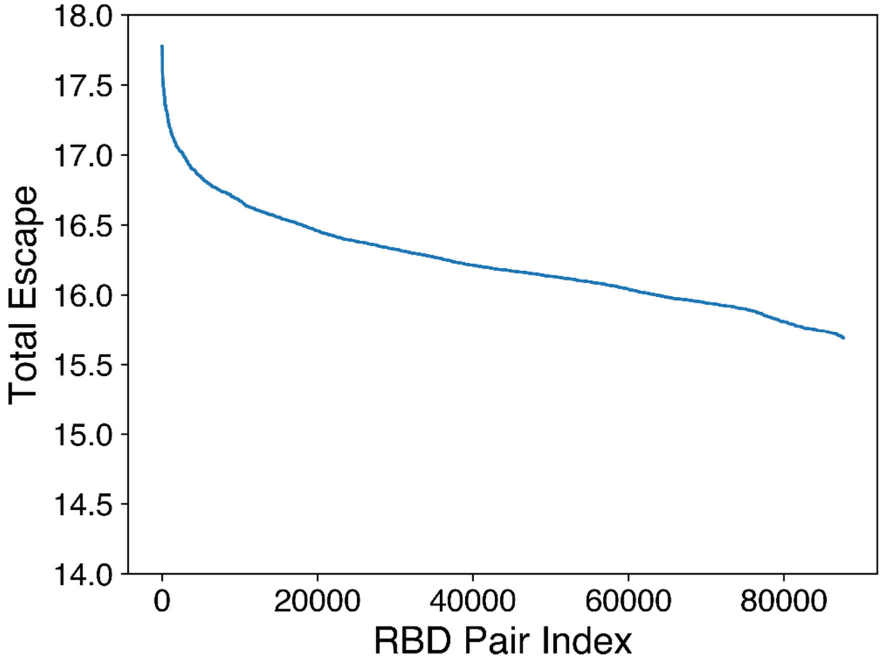


**Figure S2.** The total escape against class 1 and 2 antibodies for all ~90,000 RBD pairs that pass screening. The total escape is obtained from DMS experiments[1-5] and antibody RBD-epitope classes are defined in Barnes et al.[6]

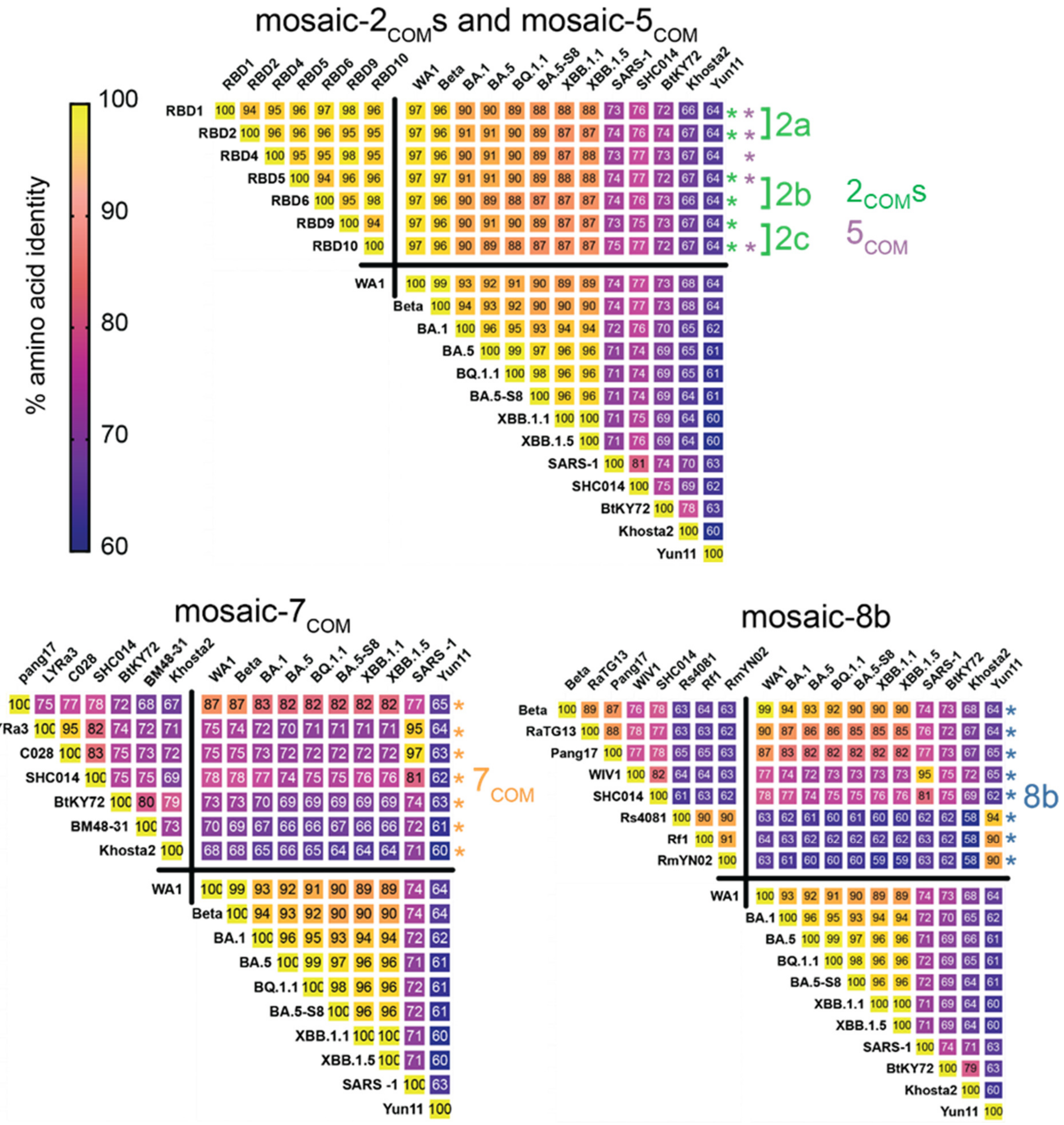


**Figure S3.** RBD amino acid sequence identities for computationally designed mosaic RBD-NPs and mosaic-8b**.** Asterisks indicate strains used for mosaic-RBD NPs.

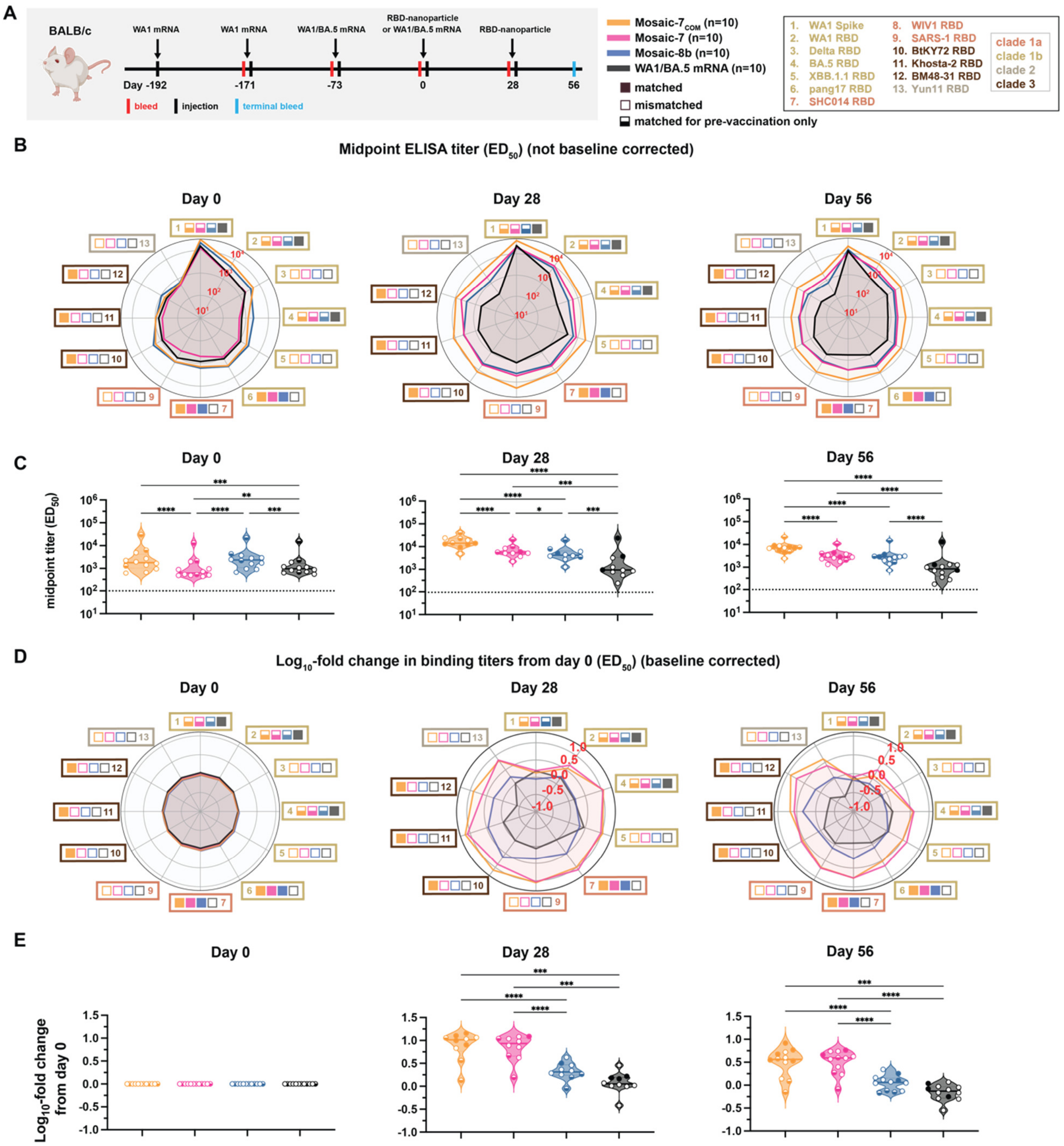


**Figure S4. Mosaic-7COM immunization in pre-vaccinated mice elicited superior cross-reactive antibody responses.** The mean of mean titers is compared in panels C and E by Tukey's multiple comparison test with the Geisser-Greenhouse correction calculated using GraphPad Prism, with pairings by viral strain. Significant differences between immunized groups linked by horizontal lines are indicated by asterisks: p<0.05 = *, p<0.01 = **, p<0.001 = ***, p<0.0001 = ****.

Binding responses at day 0 (before NP or other vaccine immunizations) showed significant differences across cohorts in titers elicited by the pre-vaccinations.[9] To account for different mean responses at day 0 between cohorts, we applied baseline corrections in Figure 6 (see Methods). Here, binding data are shown as both baseline corrected (panels B and C) and not baseline corrected (panels D and E).

(A) Left: Schematic of vaccination regimen. Mice were pre-vaccinated with mRNA-LNP encoding WA1 spike and bivalent WA1/BA.5 prior to prime and boost immunizations with RBD-NPs at day 0 and day 28 or an additional WA1/BA.5 mRNA-LNP immunization at day 0. Middle: Colors and symbols (squares) used to identify immunizations (colors) and matched (filled in), mismatched (not filled in), or matched to pre-vaccination (half-filled in) viral strains (squares). Right: numbers and colors used for sarbecovirus strains within clades throughout the figure. (B) ELISA $ED_{50}$ binding titers in serum samples from mice immunized with the indicated immunogens measured at day 0, day 28, and day 56 against spike or RBD proteins from the indicated sarbecovirus strains (numbers and color coding as in panel A). (C) Mean ELISA titers for each type of immunization at the indicated days. Each circle represents the mean $ED_{50}$ titers from mice against a single viral strain of sera from mice that were immunized with the specified immunogen (solid circles=matched; open circles=mismatched; colors for different strains defined in panel A). (D) Mean fold change in ELISA $ED_{50}$ binding titers from day 0 in serum samples from mice immunized with the indicated immunogens measured at day 0, day 28, and day 56 against spike or RBD proteins from the indicated sarbecovirus strains (numbers and color coding as in panel A). (E) Means of fold changes in ELISA titers for each type of immunization at the indicated days. Each circle represents the mean fold change in $ED_{50}$ titers from mice against a single viral strain of sera from mice that were immunized with the specified immunogen (solid circles=matched; open circles=mismatched; colors for different strains defined in panel A).
